# Compositional Recombination Relies on a Distributed Cortico-Cerebellar Network

**DOI:** 10.1101/2025.10.07.680895

**Authors:** Joshua B. Tan, Isabella F. Orlando, Jungwoo Kim, Christopher J. Cueva, Jayson Jeganathan, Giulia Baracchini, Rebekah Wong, Eli J. Müller, Claire O’Callaghan, James M. Shine

## Abstract

Human cognition depends on the ability to flexibly recombine existing knowledge in new ways. Although this capacity for compositionality has traditionally been attributed to cortical networks, its broader neural basis remains unclear. Here, we combined dimensionality reduction of task-based fMRI with recurrent neural network modelling to dissociate two processes underlying compositional cognition: the recruitment of specialised *components;* and the more general process of *recombination*. Across 87 participants performing a well-established compositional task, component processes were supported by domain-selective cortical and anterior cerebellar regions, whereas recombination engaged a distributed cortico–cerebellar network that was low-dimensional, highly integrated, and generalised across contexts. Similar functional signatures were also observed in recurrent neural networks trained to perform multiple cognitive tasks, suggesting that low-dimensional recombination is a general solution for flexible compositional cognition. Our findings revise existing models of compositional cognition by highlighting cortico-cerebellar interactions as a mechanism for flexible, integrative task generalisation.

## Introduction

Adaptive behaviour is driven by our ability to indefinitely recombine and apply existing knowledge and behaviour to novel situations or in new ways^1,2^ – a capacity described as compositional cognition^3–6^. Compositional cognition is integral in our daily lives (e.g., coming up with a new route to drive when your familiar one is blocked by road works), and for highly specialised behaviours (e.g., switching strategies in a tennis match to target your opponent’s weakness). Previous work in human functional MRI (fMRI) has typically focused on compositionality as a cerebral cortical process^3,7–11^, with a growing body of animal work highlighting the role of the prefrontal cortex and hippocampus in compositional processes^12–14^. However, an understanding of how these cortical mechanisms interact with the subcortex and cerebellum to implement compositional cognition remains incomplete.

There are two key sub-processes underlying compositional cognition: the capacity to categorise, store and recall specific *Components* (i.e., existing knowledge or behaviours), and the *Recombination* process that flexibly combines components to be applied across different contexts^6,15–17^ (Figure 1a). During compositional cognition, the two processes work in tandem: Components ensure functional specialisation within specific contexts, whereas Recombination is a flexible process that generalises across contexts^3,6,12,13,18,19^. Importantly, both Components and Recombination are essential for compositional cognition. Without a diversity of Components, behaviour will lack nuance or complexity, and without Recombination, behaviour becomes non-generalisable and yoked to each specific scenario. Here, we distinguish between the capacity to store and retrieve Components – that is, those features to which a compositional process is applied – versus the Recombination process that is typically synonymous with compositionality^5,20^. Differentiating the brain networks responsible for these distinct aspects of compositional cognition is crucial for advancing our understanding of the process.

**Figure 1.**
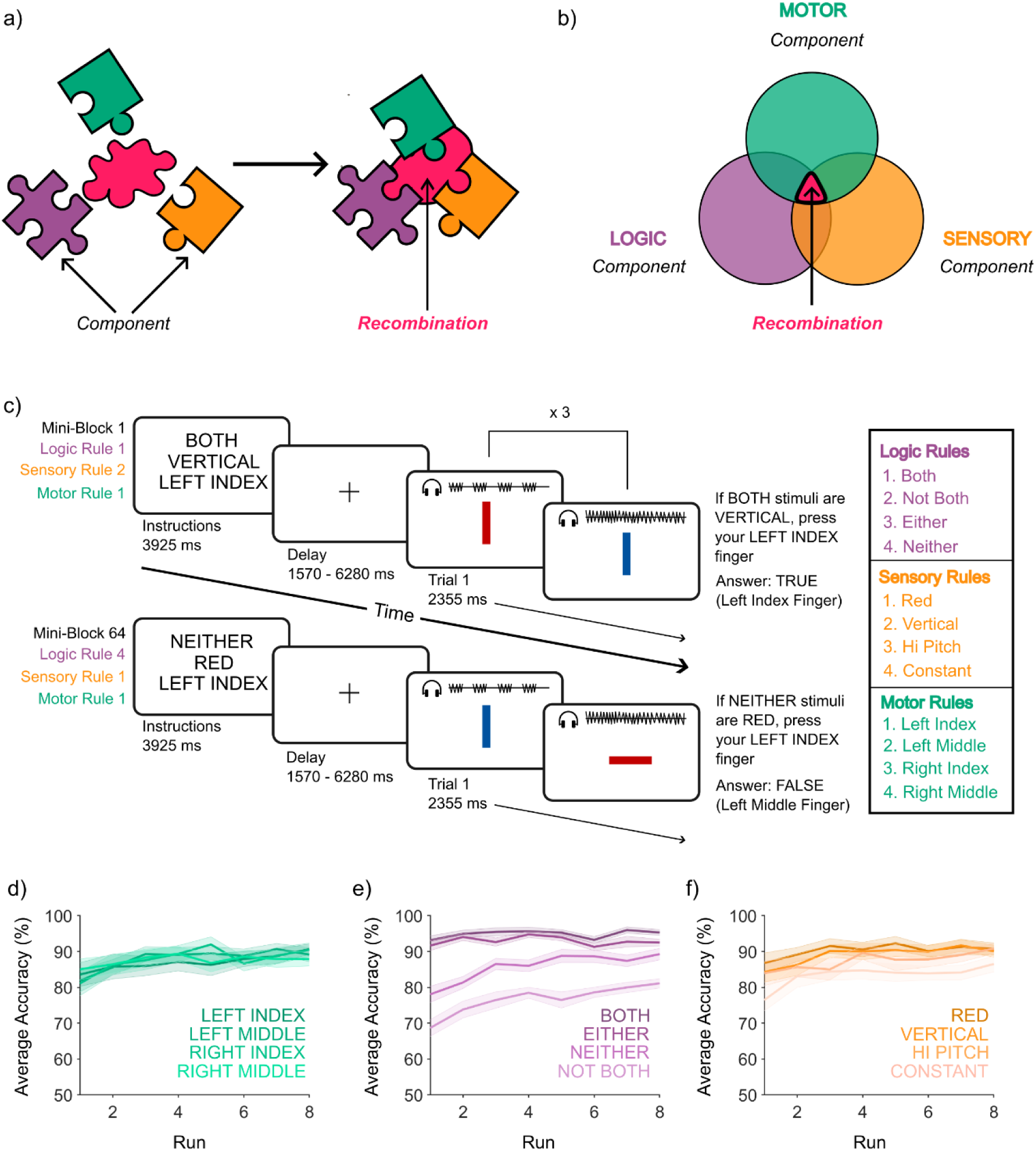
Compositional Cognition and Task Behaviour. a) A schematic of compositional processes, in which the puzzle pieces symbolise Components (different colours represent different rule domains) that can be combined by a flexible Recombination process in the centre that adapts its shape to each puzzle piece. b) the compositional process is comprised of the Recombination of different Components – each wing of the Venn diagram represents one of the context-dependent Component dimensions (Motor, Logic or Sensory), whereas the centre of the Venn diagram represents the Recombination process, which is flexible and relatively context-independent. c) the Concrete Permuted Rules Operation (C-PRO) task, wherein each trial requires the flexible Recombination of three Component domains (Motor, Logic and Sensory), adapted from^8^. d-f) Group accuracy across runs for each rule per domain d) Motor e) Logic f) Sensory. Note that mean performance was relatively stable across runs. Figures d-f were inspired by^46^.

The current leading explanations for how the human brain supports compositional cognition are predominantly cortico-centric^5,7–9,21–24^ (though see Mill and Cole^25^), neglecting a whole-brain perspective where cognitive processes are supported by interactions between distributed networks that span cortical and subcortical systems^26–29^. Among a host of important subcortical structures, a crucial system for cognitive processes is the cerebellum^30,31^. With one of the highest neuronal counts in the adult human brain, the cerebellum has previously been linked to distinct cognitive functions, such as pattern separation, skill execution, and adaptation^32–36^, all of which rely on a balance of Component separation and integrative Recombination. The cerebellum both receives and sends information to the rest of the brain^37^, including the prefrontal cortex^38^, a key area of interest identified in compositional studies^3,7,9,12,23,24^. With growing evidence that cerebellar functions contribute to both cognitive and motor processes^30^, it is possible that the cerebellum also serves a role in compositional cognition.

Linking behaviour to neural activity is possible through dimensionality reduction approaches. Prominent examples in animal literature include joint principal component analysis^39^, targeted dimensionality reduction^40^ and de-mixed principal component analysis^41^. An alternative approach that has been well-established in fMRI literature is partial least squares (PLS) analysis^42–44^, which defines a low-dimensional space that captures the maximal covariance between neural activity and task structure, allowing for the separation of different conditions and task variables. Through PLS, we identified distinct patterns relating to Component and Recombination processes.

Here, we used fMRI to analyse time-resolved BOLD responses across the cerebral cortex, subcortex, and cerebellum during performance of the Concrete Permuted Rules of Operation (C-PRO) paradigm, a task that relies on compositional cognition^8,9,45^. Our primary aims were to 1) decompose compositional cognition into Component processes and the Recombination process; 2) test the hypothesis that compositional Recombination involves distributed computations across subcortical and cerebellar structures; and 3) identify any inter-regional dynamics that may be crucial for compositionality. We hypothesised that: 1) Recombination would be consistent and generalisable across varied rule contexts, while Components would be specifically and selectively recruited for each rule; 2) Components would involve specialist regions, whereas Recombination would recruit a distributed network across cortical and cerebellar regions; and 3) networks underlying Components and Recombination should work together dynamically in order for effective compositional cognition to occur.

## Results

### Concrete Permuted Rules of Operations Task

Ninety-five participants completed the C-PRO paradigm as part of an openly available dataset (OpenNeuro, accession number ds003701)^8,9,45^. The C-PRO paradigm operationalises compositional cognition by introducing distinct domains of rules as Components that are recombined during the task (Figure 1a). By varying the specific combination of rules, the C-PRO creates different task contexts. Therefore, each trial contains information that is specific to the context (i.e., the Motor, Logic and Sensory-specific *Component* processes) and information that generalises across contexts (i.e., the *Recombination* process; Figure 1b).

Participants were presented with three rules, with one per task domain (Motor, Logic and Sensory). After a short delay, two stimuli were presented sequentially, after which participants were required to make an appropriate response. For example, if the instructions were “BOTH” (logic rule), “VERTICAL” (sensory rule), and “LEFT INDEX” (motor rule), participants would press a button with their LEFT INDEX finger if BOTH bars were VERTICAL (Figure 1c). With four possible rules per domain, there were a total of 64 unique permutations of rules. Each of these 64 task contexts were presented three times (trials) in a row, which we refer to as mini-blocks, and each mini-block was completed twice (total of 128 mini-blocks) across eight scanning runs. Participants who scored 100% on more than 64 mini-blocks were kept in the final dataset, resulting in 87 participants. For more details regarding the task paradigm refer to the *Methods*.

We fit a generalised linear mixed model that compared accuracy between rules across runs and whether there was a difference in performance between the first and second exposure to a specific mini-block, while controlling for repeated measures across runs and participants. From the model, we observed a significant difference in accuracy depending on which Logic and Sensory rules were applied, where accuracy was significantly higher for the rules “BOTH” (logic) and “RED” (sensory) compared to all others (p < 0.05; Figure 1d-f). There were no significant differences in accuracy across the different Motor rules (p > 0.05). Accuracy did not differ significantly between the first and second exposure to the mini-blocks (p > 0.05). See Supplementary Material for detailed results.

### Compositional Component Processes

To detect time-resolved task-related brain activity associated with each of the Component constructs within the C-PRO task, we subjected pre-processed BOLD data from 482 regions that covered the cerebral cortex, cerebellum, basal ganglia and thalamus^47–50^ to a finite impulse response (FIR) model (constructed across a time window of 23 TRs for each mini-block starting from trial onset). To account for different response timings across mini-blocks, the time window was specified using the maximum number of time points present across all mini-blocks, allowing us to identify signatures in the BOLD time series of compositional cognition (see *Methods*).

An inherent design feature of the C-PRO paradigm is that every trial of the task contains a blend of rules from all three domains, meaning that there is no simple means for isolating the brain patterns for each task domain. To circumvent this issue, we applied a mean-centred, task-based partial least squares (PLS) analysis^44^ to determine latent variables that capture maximal covariance between brain activity and behaviour, separating out brain patterns for each task domain. This allowed us to identify a low-dimensional orthogonal basis set associated with the task (the top three latent variables explained 88.7% of the total variance). To further confirm whether the first three latent variables were significant or occurred by chance, we created a null distribution of singular values by permuting regional brain activity 1,000 times while holding the task design matrix static. Importantly, all three latent variables were significant after permutation testing (p = 0.001), supporting the conclusion that the first three latent variables captured the maximal brain-behaviour covariance in the data explain the data. To test whether the PLS overfitted to the data, we applied a split-half test by constructing separate PLS spaces on the first (n = 44) and second halves (n = 43) of the dataset. The top three latent variables across both halves were significantly correlated (r_1_ = 0.99, p < 1e-16; r_2_ = 0.97, p < 1e-16; r_3_ = 0.87, p < 1e-16). Therefore, all further analyses were conducted in a PLS space constructed on all participants using these three latent variables.

To interpret the latent variables, we sought convergence across two distinct approaches: we mapped each C-PRO rule to the three latent PLS variables (Task Contrast Scores, Figure 2a-c) and interrogated the brain loadings on each latent variable (Figure 2d-f). Both approaches offered clearly congruent interpretations. LV_1_ was dominated by *Motor* rules (left hand vs. right hand; Figure 2a) and separated the left and right motor regions of the cerebral and cerebellar cortex (Figure 2d); LV_2_ was mainly driven by *Logic* rules (positive vs. negative; Figure 2b) and separated higher cognitive areas from unimodal and memory-associated areas (Figure 2e); and LV_3_ was driven by *Sensory* rules (visual vs. auditory; Figure 2c) and separated attentional processing and higher order visual processing from primary auditory and primary visual regions (Figure 2f). These results confirmed that the task-based PLS was able to effectively identify the three main Component processes that comprised the C-PRO task.

**Figure 2.**
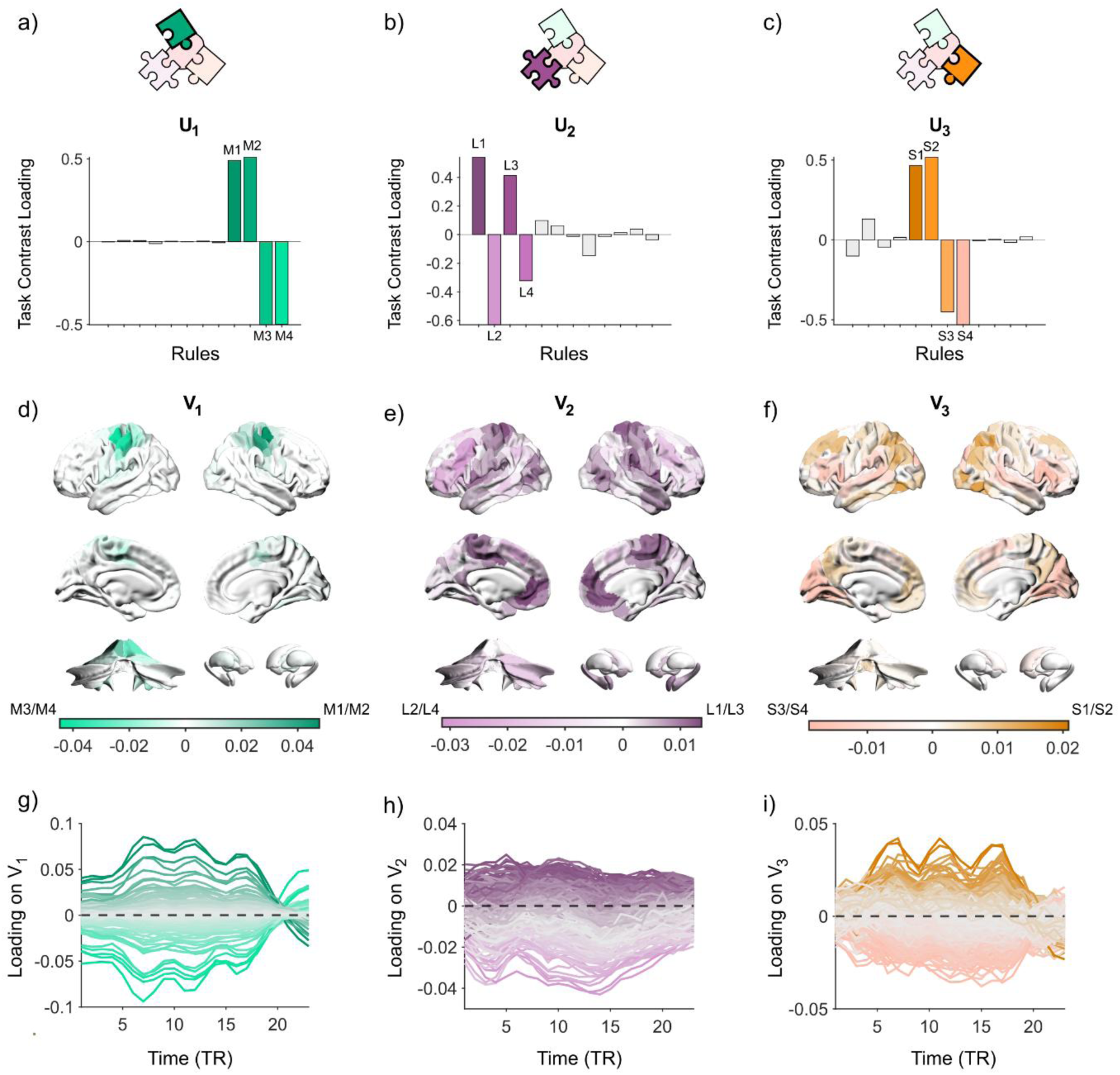
Segregated Subnetworks Are Required for Compositional Component Processes. a-c) Bar plot of the first three latent variable (LVs; U_1-3_) of Task Contrast scores in PLS space. d-f) Brain visualisation of first three LVs (V_1-3_) of average brain activity in PLS space. g-i) Loadings of brain activity across time in PLS space. The three ‘peaks’ in g) and i) relate to the three trials completed per mini-block. Each line denotes an individual region’s dynamics across time.

To track how brain region dynamics evolved in the PLS space, we plotted the region’s time-resolved loadings (Figure 2g-i). For LV_1_ and LV_3_, we observed trial-related dynamics denoted by three peaks (one per trial), whereas LV_2_ was more consistently associated with task-based negative deflections. Note that the interpretation of PLS loadings is invariant to sign, such that the negative deflections can be interpreted either as a negative loading on the positive (purple) regions of LV_2_, or as a positive loading on the negative (pink) regions of LV_2_. In this case, interrogation of the PLS loadings was consistent with the latter interpretation.

Task-related neural signatures can appear during preparation phases^51^, however it is unclear to what extent neural signatures between preparation and task execution are similar. To verify whether this decomposition was unique to the trial period or was consistent for all aspects of compositional processing, we generated a PLS space from the time series related to the task “Instruction” period. The first three LVs of the Instruction period separated the rules per domain in the same manner as the decomposition of the trial period (Supplementary Figure S1). The brain patterns for the first three latent variables in the Instruction period were similar to the brain patterns generated from the trial period (|r_1_| = 0.99; |r_2_| = 0.98, |r_3_| = 0.96, respectively; Supplementary Figure S1). A key difference between the two periods was that the regional dynamics in the PLS space for the trial period showed peaks and troughs related to each individual trial, whereas the regional dynamics of the Instruction period showed block activation dynamics (Supplementary Figure S1). Taken together, our analysis demonstrated that PLS was able to successfully separate out brain patterns and dynamics specific to each domain and these patterns were consistent and specific to compositional processing and not to task execution.

### Compositional Recombination Process

Having identified a set of axes associated with the Component processes of the C-PRO task, we next designed an approach to identify regions associated with the Recombination process. For regions to be involved in Recombination, we reasoned that: i) they should be engaged across most trials (i.e., high mean FIR response); ii) they should not be selectively associated with specific Components (i.e., PLS loading near the origin); and iii) their activity should be correlated with Component brain regions on those trials corresponding to that Component: for instance, they should co-activate with “left hand” brain regions during “left hand” trials.

We began by associating each brain region with a vector in three-dimensional space, corresponding to the first three dimensions of the PLS analysis. (Figure 3a). The absolute value of the loading on each PLS dimension is interpretable as the extent to which a particular region was associated with a specific task dimension (Figure 2). Regions that loaded towards the extremity (positive or negative) were biased towards specific rules (i.e., rule dependent Component processes), whereas regions closer to the origin (0,0) showed consistent patterns across all rules (i.e., rule independent Recombination processes) or were not recruited during the task. We quantified each region’s rule-specificity by calculating the Euclidean distance of its vector from the origin at every time point of the task.

**Figure 3.**
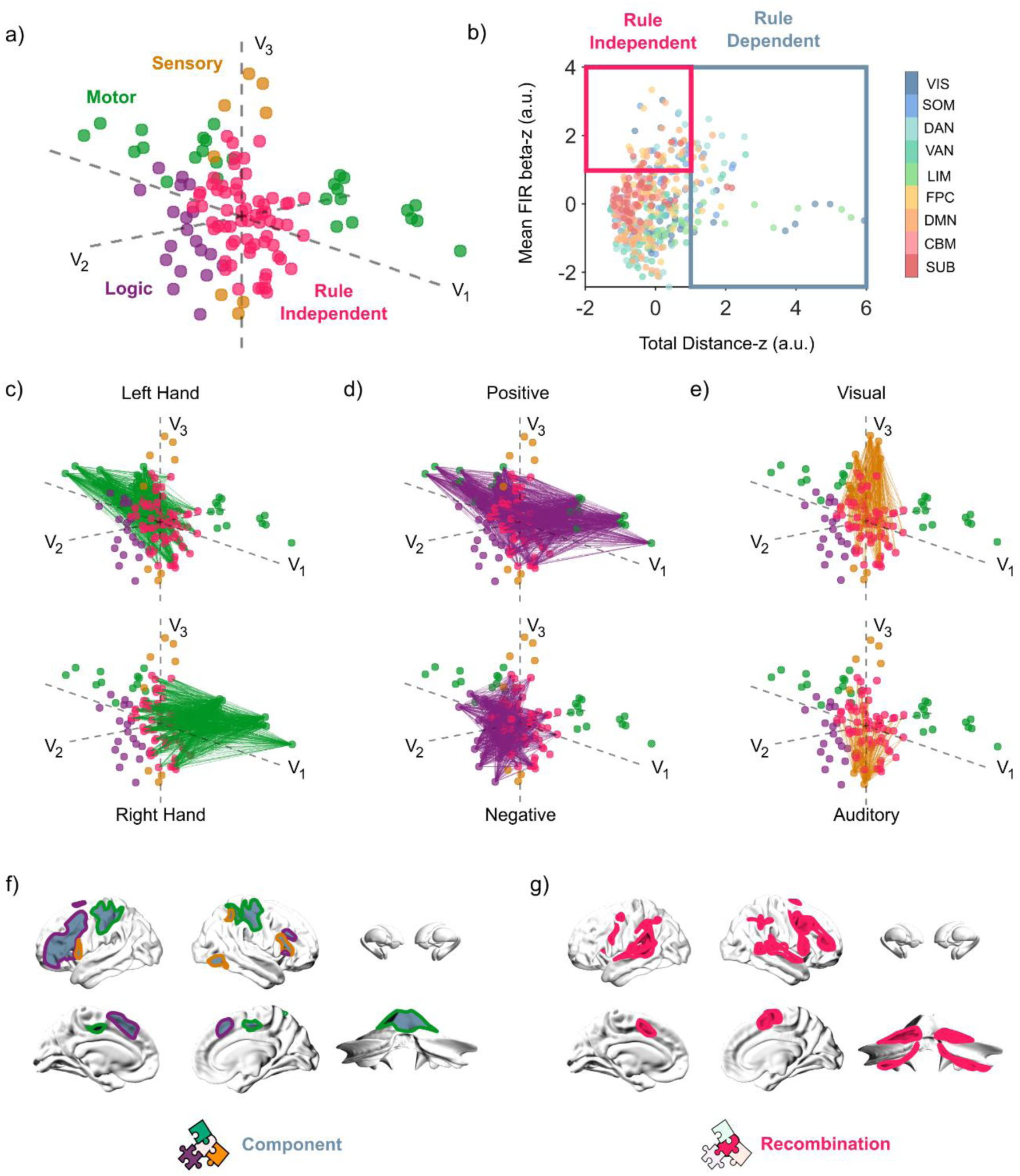
Identifying Regions Involved in Compositional Recombination. a) 3D visualisation of rule dependent and rule independent brain region using the first three latent variables when regions on the *Motor* axis were maximally separated (t = 7). Rule independent regions were coloured in pink. Rule dependent regions (green, purple, orange) were coloured by the rule domain (Motor, Logic, Sensory) that they were most biased towards. Euclidean distance was calculated for each region to the origin (0,0). b) Plotting regions by distance from the origin and average FIR estimate. Both measures were z-scored, and boundaries were drawn at x = 1 (separated rule dependent regions) and y = 1 (separated rule independent regions). Regions were coloured by the Yeo 7-networks^52^, cerebellum and subcortex. c-e) Significant functional connectivity between rule independent and rule dependent regions for each rule domain. c) Regions for Left Hand (top) and Right Hand (bottom) rules. d) Regions for Positive logic (top) and Negative logic (bottom) rules – note that Positive responses were more accurate than Negative responses (Figure 1e) and hence were more typically associated with a motor response. e) Regions for Visual (top) and Auditory (bottom) rules. f) Brain visualisation of Component regions with outlines coloured by specific rule domains. g) Brain visualisation of Recombination regions with outline.

To find the total rule bias of a given region, we measured a region’s distance from the origin across time. We then z-scored this measurement and identified rule dependent Component regions if the z-score was greater than 1 (i.e., greater than 1 *s.d.* from the mean). Using this criterion, 54/482 regions were identified as ‘rule dependent’ and were assigned labels denoting which PLS dimension they were most biased towards. These regions included the somatomotor and prefrontal cortical regions, and anterior cerebellar regions (lobules I-IV; Figure 3b, 3f, silver). From the remaining 428 regions, those that were strongly recruited during the task were identified as ‘rule independent’. Strong recruitment was measured by having a z-score > 1 from the mean FIR estimate. 65/428 regions were thus identified as ‘rule independent’ (Figure 3b, pink). These analyses identified 65 regions that fulfilled criteria *i)* and *ii)* outlined above.

To test whether the rule independent regions identified from the PLS analysis were selectively correlated with specific Components (criteria *iii*), we calculated functional connectivity matrices using Pearson’s correlation of the time series during the trial period. We compared the strength of functional connectivity patterns between rules in each domain (left hand – right hand; positive – negative; visual – auditory). Rule independent regions showed higher functional connectivity to rule dependent regions that were biased towards the specific rule – i.e., among 65 rule independent regions, 64 regions showed significantly higher functional connectivity to left hand regions in left hand contexts compared to right hand contexts (p < 0.05; Figure 3c), 56 regions showed significantly higher functional connectivity to positive Logic regions in positive Logic contexts compared to negative Logic contexts (p < 0.05; Figure 3d), and 60 regions showed significantly higher functional connectivity to visual regions in visual rule contexts compared to auditory contexts (p < 0.05; Figure 3e). Note that any bias towards positive Logic or negative Logic was driven by whether the participant responded, in which case there were significantly more incorrect responses during negative Logic contexts (Figure 1e).

A conjunction analysis across these three contexts identified 51 of 65 regions that were significant across all three Component trial types (Figure 3g). Importantly, the 51 regions we identified were distributed across both the frontoparietal cerebral cortex and the posterior cerebellum (lobules VI, VIIb, VIIIa), demonstrating a spatial dissociation in the cerebellum between Component and Recombination processes, thus providing robust evidence in support of our original hypothesis. From these sets of analyses, we have identified a set of regions serving rule dependent Component processes and a set of regions that fit our criteria for Recombination. For detailed atlas labels of Component and Recombination regions refer to Supplementary Material Tables 1 and 2.

To assess the signal reliability for Component and Recombination regions across the latent variables, we created 1,000 bootstrap samples and calculated the standard error of a region’s PLS loading across samples (refer to *Methods).* Regions with a bootstrap ratio (|BSR|) > 2 (analogous to p-value < 0.05) were defined as stable^44^. All Component and Recombination regions were stable across at least one of the three latent variables and were used in future analyses, with 51/54 Component regions and 46/51 Recombination regions stable across all three latent variables. Importantly, both anterior and posterior cerebellar regions were stable across all three latent variables (Supplementary Figure S4). To ensure that the main results were not driven by regions stable on specific latent variables, we replicated our results using a stricter threshold of |BSR| > 2 across all three latent variables. Together, our results confirm that the distributed network associated with Recombination was related to a set of task-dependent functional correlations that were specific to the Component-specific regions required to complete the C-PRO task.

### The Recombination Network is Low-dimensional, Integrated, and Generalises Across Task Contexts

The extent to which neural activity generalises has been related to the dimensionality of the neuronal time series – i.e., time series of generalisable neurons are typically low-dimensional with low separability across contexts, whereas non-generalisable neurons are more complex, high-dimensional and separable across contexts^16,36^. To apply this framework at the macro-scale, for every participant we measured the dimensionality of Recombination and Component regions separately, using the participation ratio^36,53,54^ (refer to *Methods)*. This dimensionality measure specifies the number of dimensions required to capture the variance of the data. For example, data with variance along a single dimension has a participation ratio of 1 (low-dimensional), whereas data with variance evenly spread across N orthogonal dimensions has a dimensionality of N (higher-dimensional). We computed the dimensionality of neural activity across all mini-blocks, where lower dimensionality implies a response that was shared across all mini-blocks and vice versa. We then compared the difference in dimensionality between Recombination and Component regions using a paired t-test. Component regions had significantly higher dimensionality compared to Recombination regions (t_86_ = 10.61, p < 1e-16, 95% CI [1.18 1.73]; Figure 4a), suggesting that Recombination regions were low-dimensional and shared a common response across mini-blocks, whereas Component regions were higher-dimensional with more specific responses for different mini-blocks.

**Figure 4.**
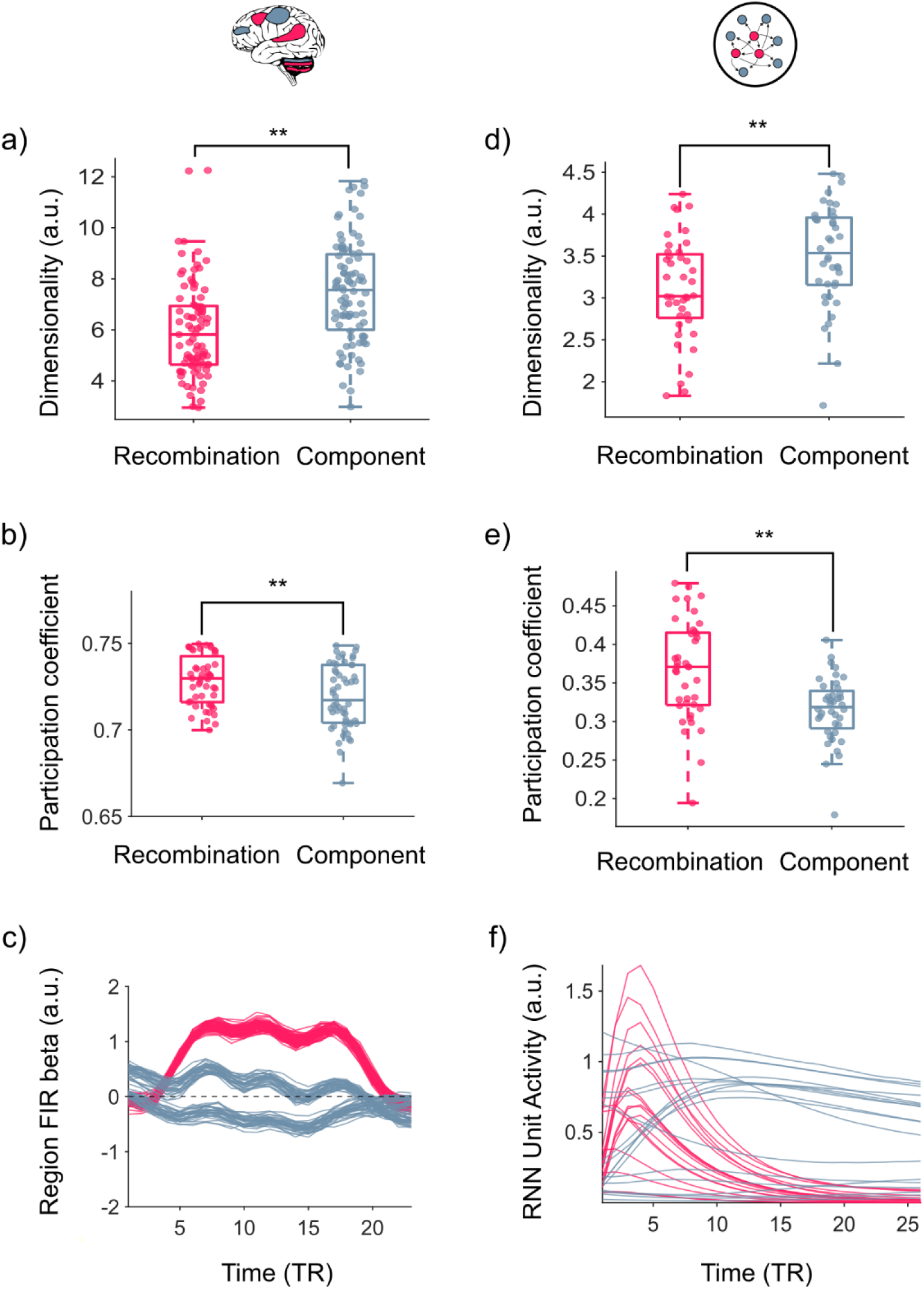
Characteristics of Recombination. a-c) Characteristics measured from the human fMRI data. a) Boxplot of Recombination (pink) and Component (silver) region dimensionality measured by the participation ratio. b) Boxplot of participation coefficient scores for Recombination (pink) and Component (silver) regions. c) Group average beta estimates across time for an example region from Recombination (pink) and Component (silver) regions. Each line is the FIR estimate for a different mini-block. d-f) Characteristics measured from simulated recurrent neural networks. d) Boxplot of Recombination (pink) and Component (silver) unit dimensionality measured by the participation ratio. e) Boxplot of average participation coefficient scores for Recombination (pink) and Component (silver) units per network. f) Average unit activity (firing rate) during the response period for an example Recombination (pink) and Component (silver) unit. For boxplots in a) and b), each dot is a brain region. Centre line, median, box limits, upper and lower quartiles. Whiskers, 1.5x interquartile range. ** denotes significant difference p < 0.01. For boxplots in d) and e), each dot indicates the Recombination (pink) or Component (silver) units within a single neural network (n = 40 networks). Centre line, median, box limits, upper and lower quartiles. Whiskers, 1.5x interquartile range. ** denotes significant difference *p* < 0.01.

A key feature of Recombination regions is their ability to flexibly interact with the rest of the brain. To measure the extent of communication between distinct brain regions, we used a measure of network integration – the participation coefficient^55^. We calculated participation coefficients for Recombination and Component regions using a population averaged functional connectivity matrix (refer to *Methods*). From these analyses, Recombination regions were found to be significantly more integrated with the rest of the brain compared to Component regions (t_103_ = 2.93, p = 0.004, 95% CI [0.003 0.016]; Figure 4b).

Both Recombination and Component regions are important for compositional cognition, however the differential contributions of these regions to the task are hard to dissociate using traditional analyses. We hypothesised that the coupling of a given region’s activation to the task context would differentiate the two groups. Using this feature, we predicted that Recombination regions would be non-specific, for example, no matter whether the mini-block uses right or left hand, the region will have the same response (Figure 4c; pink). By contrast, Component regions would be more specific to the context, for example, regions specific to right hand rules will have different patterns during mini-blocks with left hand rules (Figure 4c; silver).

To test this hypothesis, for every participant, we correlated each region’s FIR time series within each mini-block to every other mini-block. We compared the similarity of FIR estimates between Recombination and Component regions using a generalised linear mixed model, while controlling for repeated measures across participants. Consistent with our prediction, Recombination regions were more correlated than Component regions across mini-blocks (t_9,913_ = 42.58, p < 1e-16, 95% CI [0.11 0.12]). Therefore, recruitment of Recombination regions was non-specific and generalised across all mini-blocks, while recruitment of Component regions was specific to the rules being implemented. Taken together, Recombination regions were low-dimensional, integrated, and generalised across task contexts. These results were replicated using a stricter threshold of regions with |BSR| > 2 on all three latent variables (Supplementary Figure S4).

### Characteristics of the Recombination Network are shared by Compositional Neural Networks

To test whether the characteristics of the Recombination network generalised beyond the fMRI data, we analysed 40 recurrent neural networks that were trained to perform multiple cognitive tasks. Given that our goal was to probe whether low-dimensional recombination emerges as a *general principle* of flexible computation across tasks, rather than as an artefact of a single task design, we chose multi-task networks rather than training a network specifically on the C-PRO task. We reasoned that, if recombination-like signatures appear in such networks, the features we observed in the brain reflect a general computational solution, not just idiosyncrasies of the C-PRO paradigm. From the twenty possible cognitive tasks in the original study^56^, we analysed eighteen tasks that had matching temporal window lengths during the response period (see *Methods* for details).

Following a similar pipeline to the analysis of the fMRI data, for each neural network, we identified a set of units that were related to Components (specific to the task) and Recombination (generalised across tasks) processes. To quantify the task-specificity of each unit, we followed similar concepts of task variance from Yang *et al.*^56^. To see how task variance related to our previous Euclidean distance measure in the PLS space, we first applied this measure to the fMRI data. In the fMRI data, we measured variance across mini-blocks at each time point. Task variance per brain region is then calculated by averaging across time points. When comparing task variance to our original measure for rule-specificity, there was a non-linear positive relationship (Supplementary Figure S2), indicating that the two measures captured similar features in the data.

To measure task variance in the neural networks, we used the trial-averaged unit activity per task and calculated the variance across tasks for each time point. Component units were identified by selecting units that had a normalised task variance > 1. From the remaining units, units with average activity > 1 were labelled as Recombination units. Given that there was no consistency in the neural space across iterations, we identified a set of Recombination and Component units for each network separately.

Taking the Recombination and Component units, we then measured the dimensionality, integration, and correlation across tasks. Component units had a significantly higher dimensionality compared to Recombination units (t_39_ = 3.31, p = 0.002, 95% CI [0.16 0.66]; Figure 4d). Recombination units were significantly more integrated with the rest of the network (t_78_ = 5.51, p = 4.54e-07, 95% CI [0.03 0.07]; Figure 4e) and were more correlated across tasks compared to Component units (t_1,255_ = 8.46, p < 1e-16, 95% CI [0.12 0.19]; Figure 4f). Overall, the results from the neural networks aligned with our findings from the fMRI data, demonstrating that these characteristics are underlying features that separate Recombination and Component processes during compositional cognition, irrespective of the specific implementation-level details of each system.

## Discussion

Compositional cognition can be characterised by two related, yet separable, processes: the specialised Components that are recruited to achieve the task demands, and the more generalised process of Recombination that facilitates these components to be applied in novel situations. Here, we used dimensionality reduction, time-varying modelling and functional connectivity to interrogate fMRI measurements collected during a compositional task (C-PRO) to dissociate these processes. We found clear evidence that the two processes differed. First, Components were embedded in specific contexts and rules, predominately identifiable in specialist cortical regions, whereas Recombination was a feature that generalised across all contexts, involving a distributed cortico-cerebellar network. We demonstrated that compositional cognition required specific, dynamic interactions and differentiation between these two processes, and that the Recombination network was low-dimensional, integrated, and generalised across task contexts. Finally, we observed that the network signatures of Recombination observed in human neuroimaging data were also present in recurrent neural networks trained on multiple cognitive tasks. Overall, these results provide evidence that distributed network interactions underpin two distinct processes forming compositional cognition.

During compositional cognition, generalisation is important for flexibility and applying similar behaviours across many different contexts, whereas specialisation is important for distinguishing between behaviours and adding complexity and nuance^16,19^. In human neuroimaging studies, brain networks have been described using specialised or generalised functions. For example, task-based fMRI studies have described a range of brain networks from specialised sensory functions (i.e., motor network)^57^ to generalised higher cognitive functions (i.e., cognitive control)^58^. However, typically these networks encompass the entire task – e.g., generalised networks are recruited during tasks that recruit higher cognitive functions, such as the n-back task. Our study expands on these frameworks, applying measurements from neural population studies^59,60^ to demonstrate that both specialised and generalised brain networks exist during compositional cognition. Our results identified two distinct task-related networks consisting of Component and Recombination regions. Component regions were tightly coupled to specific rules and contexts – e.g., regions recruited for right-hand movements were not recruited during left-hand movements, whereas Recombination regions were recruited regardless of the context and rule. By characterising macro-scale signatures of task fMRI through neural population measures, we offer a nuanced whole-brain perspective on how specialised and generalised networks give rise to compositional cognition.

Recombination recruited a distributed cortico-cerebellar network that was not associated with specific rules of the paradigm. Recombination is typically associated with specific brain areas, such as the frontoparietal control network^7,9^ and hippocampus^5,14^. Our results expand on this framework, identifying a distributed cortico-cerebellar network important for recombination. Crucially, while some rule independent regions overlapped with frontoparietal areas, we also found evidence for regions from the superior temporal cortex/temporoparietal junction and cerebellum to serve a role in Recombination. We found no significant results for the thalamus and basal ganglia, possibly due to the lower signal-to-noise ratio and the small size of the nuclei^61^. The superior temporal cortex/temporoparietal junction have been related to highly integrative cognitive capacities, including language and social processing/theory of mind^62,63^. Capacities that are synonymous with an ability to apply a set of known rules or components (e.g., syntax and grammar; or social norms and conventions) in novel ways and across novel contexts. The C-PRO exemplifies the re-use and recombination of rules across contexts, echoing the combinatorial processes that form the basis of language and complex social interactions^64^.

There is growing evidence that the cerebellum which is typically associated with motor movements and coordination is also important for cognitive processes such as pattern separation and prediction^30,35,65^. From our results, we dissociated between anterior cerebellar lobules (I-V) and more posterior lobules (VI, VIIb, and VIIIa), where the anterior lobules were related to Components and rule specialisation and the posterior lobules were related to Recombination and generalisation. This dissociation follows with overall cerebellar functional patterns that dissociate between motor regions (anterior) and higher order executive functions (posterior)^66^. Lobules VI, VIIb, and VIIIa have specifically been associated with motor planning^67^, and the multi-demand network^68^, including functions such as updating, shifting, and inhibition^69,70^.

During recombination, depending on how components are recombined, there will be different behavioural outcomes. Due to the cerebellum’s role in pattern separation^35^, the cerebellum can aid in developing internal models that control how components are recombined during compositional cognition^33,71,72^. Through these models, the cerebellum contributes to refining the recombination of task-components. In cats and monkeys, the cerebellum shares anatomical connections with the temporal cortex^73^, therefore it is possible that these connections were also preserved in the human brain therefore forming a cortico-cerebellar circuitry specifically recruited for recombination. To gain a more concrete understanding of this cortico-cerebellar circuitry future studies can try manipulating outputs from the cerebellum^36^ while recording from the temporal cortex. Hence, contributions from the cerebellum to the temporal cortex may facilitate recombination and generalisation of actions across contexts.

Task networks dynamically interact depending on the task context. In compositional cognition and generally across task fMRI studies, connectivity approaches typically analyse the whole task, identifying a consensus task network – through averaging and thresholding – that is “on” during the task and “off” in other stages. These approaches typically lack the nuance to identify networks that underlie the cognitive function of interest versus networks that result from the task design. For example, motor responses are useful in signalling whether a participant completes the task, however this can result in motor networks appearing in task fMRI results including higher cognitive functions^74^. To account for influences of task design on BOLD signatures, studies have explored modelling^75^ and regressing out task responses and timing^76^. While regressing out task responses accounts for task design biases, it can also remove context specific dynamics between brain networks. In this study, we followed an alternative approach by first separating out networks related to the cognitive function and networks related to task design. Using this approach, we revealed that a generalised recombination network flexibly interacts with a specialised component network, increasing connectivity to regions depending on the task context. These results demonstrate that both networks are important for compositional cognition and dynamically interact through the task. Understanding interactions between component and recombination networks provides a dynamic perspective on how brain networks meet task demands.

Developing a paradigm that can elicit compositional cognition in combination with human fMRI is non-trivial. Compositional task designs require components that are easily learned and able to be combined into complex paradigms, with enough variation to induce novelty throughout the task. Among the paradigms that have achieved this balance^3,5,8^, the C-PRO paradigm stands out as a well-established example^8,9,45^, with recent updates to the C-PRO paradigm further improving the ecological validity of the task stimuli^25^. However, one aspect that is lacking in the C-PRO paradigm is the range of task variability. During the C-PRO paradigm, task variations occur on the scale of 1-3 rules. While this is optimal for adaptation across explicitly similar skills (e.g., adapting tennis skills to pickle ball), adaptation can also occur between skills that overlap at an abstract level (e.g., adapting tennis skills to chess)^77^. Currently, adaptation at the abstract level has been explored in neural networks^56,78^ but is yet to be explored in human fMRI. We showed that Recombination features in the fMRI data were identifiable in compositional neural networks. Future studies could explore compositional cognition at the abstract level by having individuals perform a multitude of tasks that overlap at an abstract level.

Compositional cognition is facilitated by two distinct processes: Recombination and Component engagement. Between these two processes, there are distinct features. The Recombination network recruited a cortico-cerebellar circuitry that was low-dimensional, integrated and generalised across task contexts. Whereas task components were relatively high-dimensional, less integrated, and were recruited during specific task contexts. These characteristics were reproduced in compositional neural networks trained on multiple cognitive tasks. Together, these findings demonstrate that compositional cognition is facilitated by interactions between distributed networks serving Recombination and Component processes.

## Methods

### Experimental Design

Ninety-five healthy adults (mean age = 22.23 years, s.d. = 3.93; female:male = 54:41; right-handed) were recruited from Rutgers University and the surrounding Newark, New Jersey community. All participants completed the C-PRO paradigm while undergoing simultaneous MRI.

The C-PRO paradigm was designed to test rapid instructed task learning and adaptive behaviour by recombining various task rules^4,8^. We used the modified version introduced by Ito *et al.*^8^. This version utilised 12 task rules that were categorised under three domains: four logic rules (BOTH, NOT BOTH, EITHER, NEITHER), four sensory rules (RED, VERTICAL orientation, HIGH PITCH sound, CONSTANT tone), and four motor rules (LEFT INDEX, LEFT MIDDLE, RIGHT INDEX, and RIGHT MIDDLE finger). During a trial, a single rule from each domain is combined (e.g., BOTH, RED, LEFT INDEX). By matching every rule with every other rule across all domains a total of 64 unique permutations were created.

During the task, participants completed three trials in a row for each of the 64 unique permutations which we called a “mini-block”. Each mini-block lasted a total of 28,260 ms (36 TRs) and consisted of an instruction period of 3925 ms (5 TRs), followed by a jittered delay between 1570 – 6280 ms (2 – 8 TRs) sampled from uniform distribution, then three trials, where each trial is 2355 ms (3 TRs) with an inter-trial interval of 1570 ms (2 TRs), followed by another jittered delay between 7850 – 12,560 ms (10 – 16 TRs, randomised) before the start of the next mini-block. For each trial, participants were presented two sets of simultaneous audiovisual stimuli (Figure 1c). The audio stimulus was either a high pitch sound or constant tone, and the visual stimulus was a bar either in vertical or horizontal orientation, and either blue or red coloured. Response time and accuracy for each trial was measured. The paradigm was presented using E-Prime software version 2.0.10.535^79^.

Participants completed all 64 mini-blocks twice spread across eight runs (16 per run). Within the first four runs, participants completed all 64 unique mini-blocks before completing all mini-blocks for a second time in the last four runs. Order of mini-blocks was randomised between participants and runs. Alongside the eight task scanning runs, each participant also completed a resting-state scan however for the purpose of this study, only the task runs were analysed. Before entering the scanner, participants were trained on four of the possible 64 mini-blocks. The four trained mini-blocks were chosen such that all 12 rules were equally practiced.

### Imaging Acquisition

Data were collected at the Rutgers University Brain Imaging Center (RUBIC). Whole-brain multiband echo-planar imaging (EPI) acquisitions were collected with a 32-channel head coil on a 3T Siemens Trio MRI scanner with TR = 785 ms, TE = 34.8 ms, flip angle = 55°, Bandwidth 1924/Hz/Px, in-plane FoV read = 208 mm, 72 slices, 2.0 mm isotropic voxels, with a multiband acceleration factor of 8. Whole-brain high-resolution T1-weighted and T2-weighted anatomical scans were also collected with 0.8 mm isotropic voxels. Spin echo field maps were collected in both the anterior to posterior direction and the posterior to anterior direction in accordance with the Human Connectome Project preprocessing pipeline ^80^. A resting-state scan was collected for 14 mins (1070 TRs), prior to the task scans. Eight task scans (runs) were subsequently collected, each spanning 7 min and 36 s (581 TRs). Each of the eight task runs (in addition to all other MRI data) were collected consecutively with short breaks in between (participants did not leave the scanner).

### Pre-processing

Neuroimaging data was organized in BIDS format and pre-processed with fMRIprep version 23.1.4 (https://fmriprep.org/en/stable/<u>#</u>), a standard pipeline that incorporates toolboxes from the gold-standard preprocessing software in the field^81^. fMRIPrep involves the basic pre-processing steps (co-registration, normalization, unwarping, noise component extraction, segmentation, skull-stripping, etc.) and produces a report for quality checking at each step. See Supplementary Material for a full description of each step. Regression of twelve head motion artifacts (translation and rotation estimates plus the first derivatives), and the average combined white matter, CSF signal was conducted using custom python scripts, with a high-pass filter set at 0.01 Hz. This approach effectively reduces head motion artifacts while preserving task-related signal. Using custom python script, mean BOLD time series were extracted from 400 Schaefer cortical regions - a parcellation that is homogenous across both task and resting-state fMRI protocols)^48^, 28 cerebellar regions from a spatially unbiased atlas that preserves the anatomical detail of cerebellar lobules^47,50^, and 54 subcortical regions from a high-resolution atlas that accounts for individual variation^49^. These atlases were chosen as robust and popular parcellations that reveal meaningful neurobiological features^47–50,52^.

### Behavioural Analysis

Accuracy was calculated as the percentage of correct responses within a mini-block. We thresholded performance by the number of mini-blocks participants got 100% (3/3) correct. Participants that performed at 100% accuracy across more than 64 mini-blocks were kept for further analyses, leaving 87 participants in the final dataset.

Behavioural data was fit using a generalised linear mixed model, comparing the effects of different rules and learning over time against accuracy in a mini-block. Fixed effects were measured for each domain (logic, sensory, motor) separately, accuracy across runs, and whether it was the first or second time completing the mini-block in the scanner (instance) (Equation 1). Each domain was treated as a categorical variable (rules 1 – 4). Random effects included repeated measures per run and participant.

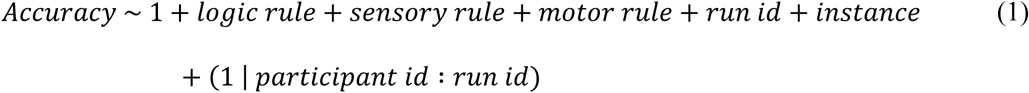

### Finite Impulse Response Model (FIR)

The finite impulse response (FIR) model estimates changes in BOLD activity at multiple time points and is not biased towards a particular shape, such as the hemodynamic response function which is typically used in task-based functional MRI analyses^82^. To choose an appropriate window of time, we measured the maximum number of time points available across all mini-blocks. Due to some mini-blocks occurring towards the end of the scan, we chose 23 time points or TRs (∼18 seconds), starting from trial onset. Each mini-block (n = 64) was modelled separately in the FIR model, resulting in 23 estimates per mini-block. Random effects for the run number and instance were included, allowing intercepts to account for repeated measures. The final design matrix had 23 regressors per mini-block (fixed effects), and regressors for which run and which instance (random effects; Equation 2). The FIR models were fit at the participant-level using generalised linear mixed models to compare the changes in the design matrix variables with a participant’s BOLD time series concatenated across runs (Equation 2). By concatenating time points within each mini-block, temporal blurring between adjacent trials was minimised and common effects across similar mini-blocks were emphasised.

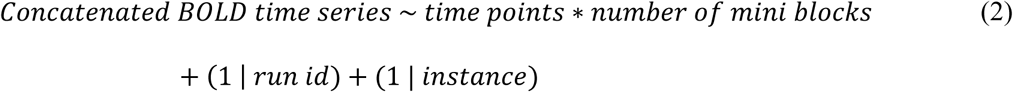

### Partial Least Squares Analysis

Partial least squares (PLS) analysis is a supervised multivariate approach that finds the maximal covariance between two sets of variables, oftentimes applied in the context of brain-behaviour relationships. The two main components of PLS are the X matrix that consists of brain activity, and the Y matrix which consists of behavioural data. The PLS method used here follows from mean-centred task PLS correlation as described by Krishnan *et al.*^44^. For the X matrix, we used the beta estimates from the FIR model. The beta estimates were concatenated across participants resulting in number of rows equal to number of participants (87) × number of mini-blocks (64), and number of columns equal to number of time points (23) × number of regions (482). For the Y matrix, we dummy coded in task contrasts specifying labels for each rule separately. This resulted in a matrix where rows equals the number of participants (87) × number of mini-blocks (64), and the number columns equals the number of rules (12).

To compare the X and Y matrices, we calculated the covariance matrix M (Equation 3).

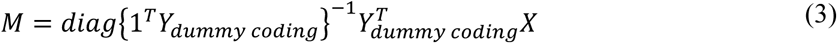

{1^T^Y_dummy coding_} is the sum of each column in the Y matrix. Diag{} is the diagonal values from the matrix. Y^T^ is the transpose (swap rows and columns) of the Y matrix.

The covariance matrix M was then mean-centred for each column by subtracting the mean of the column from all values within the column. To find the latent variables of this matrix, the demeaned covariance matrix M_mean-centreed_ underwent singular value decomposition (SVD). This resulted in three matrices: U, which is a left singular vector (latent variables) that characterises axes of behaviour in PLS space; S, singular values (diagonal matrix); and V, the right singular vector (latent variables) characterising axes of brain activity in PLS space. This approach optimally separates each task rule, and shares similarities with discriminant analysis approaches^83,84^

To decide how many latent variables to include we calculate the explained variance (Equation 4), and cumulative explained variance using the singular values from S.

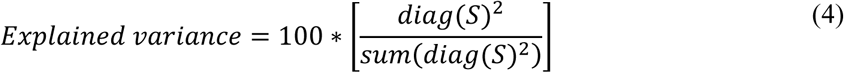

By plotting explained variance using a scree plot, we can observe how many latent variables were included before added explained variance asymptotes (elbow)^85^. From looking at explained variance, and the cumulative explained variance, the first three latent variables were used to characterise the PLS space. In this PLS space, each latent variable forms an axes that contrasts different groups of task rules.

To verify the latent variables, we used permutation testing^42,43^. Rows of the X matrix were randomly shuffled 1000 times, with Y matrix remaining unchanged. The same steps applied for each shuffled X matrix, such as calculating the covariance matrix of X_shuffled_ and Y, followed by SVD. The singular values from S were used to create a null distribution for each latent variable. The original singular values were then compared against the null distribution.

To measure the reliability of PLS loadings, we created bootstrap samples by randomly sampling with replacement. To account for repeated measures of participants, we randomly sampled participants 1000 times. Then, corresponding rows of the X and Y matrix for the sampled participant were used to create the bootstrap sample. The same steps were applied for each bootstrap sample, such as calculating the covariance matrix, followed by SVD. We calculated the standard error of each region’s PLS loading across the bootstrap samples. The bootstrap ratio was calculated by dividing the original region PLS loadings by the standard error.

To check whether the PLS analyses was not the result of overfitting, we split the data in half (n = 44 and n = 43 participants). The PLS space was generated for both halves separately and we compared the relationship between the top three latent variables of matrix V between the two halves using Pearson’s correlation.

To validate that the patterns we found from PLS were stable across both task execution and preparation phases, we applied the PLS analysis to time points during the instruction phase. To compare between the trial and instruction phases, we calculated the Pearson’s correlation for the corresponding top 3 latent variables of matrix V.

### Identifying Component and Recombination Regions

Characterising regions as rule dependent Components was dependent on a region’s bias to specific rules (measured using Euclidean distance) in the PLS space. Identifying regions involved in Recombination was based on the following criteria: i) they should be engaged across many different trials of the task; ii) they should not be selectively associated with particular task rules (Components); and iii) they should show correlated activity with task-specific dimensions on relevant task trials.

Using regional loadings in the first three latent variables from matrix V in PLS space, we created a 3-dimensional co-ordinate space for regions across time, where extreme values suggest bias towards rules, and values closer to the origin (0,0) suggest no bias. To measure regional rule bias across time, we calculated the Euclidean distance of a region from the origin at each time point. For each region, we calculated the total sum of their rule bias across time and normalised the data by z-scoring. Regions that had Euclidean distance z-score values > 1 were categorised as integral for Component (rule dependent) processes. For the remaining regions, we calculated the group average FIR time series using estimates from the FIR model. This average estimate was then normalised by z-scoring. Regions with z-scored values > 1 were categorised as rule independent. Component regions were assigned to specific domains (Motor, Logic, Sensory) depending on which PLS axes the region had the most extreme value on (e.g., regions with large positive/negative loadings on LV_1_ were assigned to the motor domain). These labels were used for the functional connectivity analyses.

### Functional Connectivity Between Components and Rule Independent Regions

Functional connectivity matrices were generated by calculating the Pearson’s correlation between rule independent and rule dependent Component regions’ time series during the trial period. Average functional connectivity matrices were calculated for each contrast as identified from the PLS analysis (left hand, right hand, positive, negative, visual, auditory rules). These matrices were vectorized and pairwise pattern similarity was estimated using Pearson’s correlation between all possible pairs.

Delta matrices between contrast pairs (left hand – right hand, positive – negative, visual – auditory) were calculated by comparing functional connectivity matrices for each contrast (Supplementary Figure S3). Rule dependent Component regions (columns) were rearranged into their assigned domains from the PLS analyses. We next ran a set of two-sample t-tests comparing different axes of each Component process (p < 0.05) and then constructed a conjunction analysis by identifying Recombination regions that showed significantly different regional functional connectivity across all three comparisons.

### Characteristics of Recombination Regions

To measure the dimensionality of regions, we concatenated the group averaged FIR time series across all mini-blocks for Component and Recombination regions and calculated principal component analysis (PCA) on each group separately. We then measured the dimensionality using the following equation^54,86^:

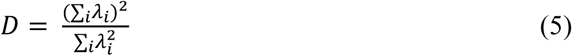

Where λ_i_ is the explained variance for the *i*th principal component. Equation 5 is commonly called the “participation ratio” and quantifies the effective number of principal components needed to explain the data. Higher values of D indicate higher dimensionality. Intuitively, when the variance of the data has equal variance along N principal components then the dimensionality is D = N. Note that in the usual scenario where the variance is not equally distributed among multiple principal components, the dimensionality will be a continuous number versus an integer. A paired t-test was used to compare the dimensionality of Component and Recombination regions.

To measure the extent a region was integrated with the rest of the brain, we followed standard pipelines from the Brain Connectivity Toolbox^55^. We calculated a population average functional connectivity matrix from trial period time series. Then, we identified modules using the Louvain algorithm. We performed a sweep across gamma values 0.5 to 2 with increments of 0.1. At each step we ran 100 iterations and calculated the adjusted mutual information^87–89^. Stability was assessed by measuring the standard deviation of modularity (Q) at each gamma value. There was low variability across all gamma values with gamma = 1 having the highest adjusted mutual information. Participation coefficient was calculated for Recombination and Component regions using the module assignment from gamma = 1. A two-sample t-test was used to compare participation coefficient between Recombination and Component regions.

Similarity across mini-blocks was measured by taking the group average FIR time series and comparing how similar (Pearson’s correlation) were time series of a given mini-block (e.g., mini-block 1) and every other mini-block. This was done for both rule independent and rule dependent regions at the participant level. To compare whether rule independent regions generalised across task contexts more than rule dependent regions, we averaged across similarity comparisons for both groups of regions at the participant level. Then we used a generalised linear mixed model to compare the average similarity value between the two groups (Equation 6).

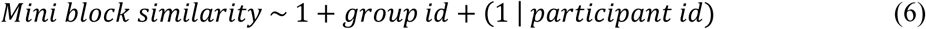

### Recurrent Neural Network Structure

Previously, Yang and colleagues^56^, trained 40 recurrent neural networks to perform across 20 cognitive tasks. Each network consisted of 256 units, which followed the following continuous dynamical equation.

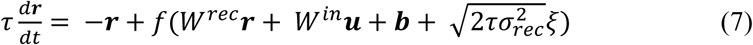

In this equation, τ = 100 ms is the neuronal time constant. **u** is the input to the network, **b** is the bias or background input, *f* (⋅) is the neuronal nonlinearity, ξ are N_rec_ independent Gaussian white noise processes with zero mean and unit variance and σ_rec_ = 0.05 is the strength of the noise. In the reference setting, we used a standard Softplus function. Each network was trained with all the tasks randomly interleaved with equal probabilities. For more details regarding the network structure and training refer to Yang *et al.*^56^.

From the 20 cognitive tasks, we analysed unit activity from 18 tasks. These included Go, React (RT) Go, Delay (Dly) Go, Anti Go, Anti RT Go, Anti Dly Go, Decision-making (DM) 1 and 2, Context DM 1 and 2, Multisensory DM, Delay DM 1 and 2, Multisensory Delay DM, and four Matching tasks (DMS, DNMS, DMC, DNMC). For DMS and DNMS tasks, two stimuli were matched if they pointed towards the same direction. In DMC and DNMC tasks, two stimuli were matched if their direction belonged to the same category, where the first category ranged from 0° to 180°, and the second category ranged from 180 to 360°. The 18 tasks were chosen due to matching number of time points during the response phase of each task. Unit activity was simulated using the continuous dynamical equation for the equivalent of 1000 trials for each task. For the rest of the analyses, only unit activity during the response phase was analysed.

### Characterising Component and Recombination processes in the Neural Networks

Yang *et al.*^56^ measured the task specificity of a neural network unit using task variance. We adapted this measure to the fMRI data by measuring at each time point, the variance of FIR time series across mini-blocks. The task variance for every region by averaging the variance across time points. We then plotted the task variance against the region’s rule bias (Euclidean distance in PLS space).

In the neural networks, we measured the task variance of each unit by taking the average unit activity per task and calculating the variance across tasks at each time point. The task variance was then measured as the average variance across time points for each unit. Units with a z-scored task variance > 1 were identified as Component units. For the remaining units, we then measured the average unit activity across all tasks and time points. Recombination units were identified with a z-scored average activity > 1. This process was completed for each neural network separately.

For each neural network, we measured the dimensionality, integration, and task similarity. Dimensionality was measured using the same previous equation but calculated across tasks instead of mini-blocks. Dimensionality of Component and Recombination units were compared using a paired t-test.

To measure integration, a correlation matrix was generated for each neural network. For each network, we ran the Louvain algorithm for 100 iterations at gamma = 1 (matching the fMRI data). An agreement matrix was constructed across iterations and a consensus partition was generated for each neural network. Across neural networks, only one network had four modules, while all other networks had five modules. For each network, the participation coefficient was calculated for Component and Recombination units. To adjust for differing number of Component and Recombination units across networks, we calculated a network average participation coefficient for Component and Recombination units. A generalised linear mixed model was then used to compare the network average participation coefficient between Component and Recombination units, and controlled for repeated measures from the same neural network.

To measure task similarity, we correlated the average unit activity for a task against all other tasks. We then compared the task similarity of Component and Recombination units using a generalised linear mixed model that controlled for repeated measures from the same neural network.

## Acknowledgements

The authors acknowledge the University of Sydney HPC service at The University of Sydney for providing HPC resources that have contributed to the processing of the data contained within this data collection.

## Funding

Australian National Health and Medical Research Council Fellowship 2016866 (CO)

Australian Research Council DP240101295 (JMS)

Australian Research Council DP250102186 (JMS, CO, EJM)

Canadian Institutes of Health Research MFE193920 (GB)

Discovery Early Career Researcher Award DE250100540 (EJM)

National Health and Medical Research Council GNT1193857 (JMS)

Research Training Program Stipend SC3227 (JBT)

University of Sydney Robinson Fellowship (CO)

## Author Contributions

Conceptualization: JBT, JMS, EM, CO

Methodology: JBT, IFO, RW, JK, JJ, GB, EM, JMS, CO, CC

Formal Analysis: JBT Visualization: JBT

Writing—original draft: JBT, JMS, CO

Writing—review & editing: JBT, IFO, RW, JK, JJ, GB, EM, JMS, CO, CC

All roles and responsibilities of the co-authors were agreed on.

## Data and Code Availability

All imaging and behavioural data are publicly available in an OpenNeuro repository (accession number ds003701). Analysis of both the behavioural and functional MRI data was conducted in MATLAB v2022b. Code required to reproduce the statistical analyses and figures are publicly available at https://github.com/ShineLabUSYD/Compositionality_C-PRO.

