## Supplementary Materials for "Compositional Recombination Relies on a Distributed Cortico-Cerebellar Network"

**1. Behaviour Statistics**

**2. fMRIprep preprocessing**

**3. Partial Least Squares analyses on Instruction period time series**

**4. Comparing Task Variance vs. Rule Bias in fMRI**

**5. Table 1. Labels for Component brain regions**

**6. Table 2. Labels for Recombination brain regions**

**7. Delta Functional Connectivity Matrices**

**8. Recombination and Component characteristics with strict Bootstrap threshold**

**1. Behaviour Statistics**

For the Logic rules, accuracy was significantly higher for the rule "BOTH" compared to all others (BOTH vs. NOT BOTH: $t_{11,124} = -28.87$, 95% CI [-0.19 -0.17], $p < 1e-16$; BOTH vs. EITHER: $t_{11,124} = -3.15$, 95% CI [-0.03 -0.01], $p = 0.0016$; BOTH vs. NEITHER: $t_{11,124} = -14.47$, 95% CI [-0.10 -0.08], $p < 1e-16$; Figure 1d). For the Sensory rules, accuracy was significantly higher for the rule "RED" compared to the auditory rules "HI PITCH" and "CONSTANT" (RED vs. HI PITCH: $t_{11,124} = -4.27$, 95% CI [-0.04 -0.01], $p = 1.98e-05$; RED vs. CONSTANT: $t_{11,124} = -11.03$, 95% CI [-0.08 -0.06], $p < 1e-16$; Figure 1e). Accuracy increased by 1% across runs ($t_{11,124} = 3.61$, 95% CI [0.01 0.02], $p = 3.08e-04$). There were no significant differences in accuracy across the different Motor rules, or whether it was the first or second exposure to the mini-block ($p > 0.05$). In summary, different Logic and Sensory rules influenced overall accuracy during the task with participants slightly improving in accuracy across runs.

### 2. fMRIprep preprocessing

#### Anatomical data preprocessing

A total of 1 T1-weighted (T1w) images were found within the input BIDS dataset. The T1-weighted (T1w) image was corrected for intensity non-uniformity (INU) with `N4BiasFieldCorrection` [@n4], distributed with ANTs 2.3.1 [@ants, RRID:SCR_004757], and used as T1w-reference throughout the workflow. The T1w-reference was then skull-stripped with a *Nipype* implementation of the `antsBrainExtraction.sh` workflow (from ANTs), using OASIS30ANTs as target template.

Brain tissue segmentation of cerebrospinal fluid (CSF), white-matter (WM) and gray-matter (GM) was performed on the brain-extracted T1w using `fast` [FSL 6.0.3:b862cdd5, RRID:SCR_002823, @fsl_fast]. Volume-based spatial normalization to two standard spaces (MNI152NLin6Asym, MNI152NLin2009cAsym) was performed through nonlinear registration with `antsRegistration` (ANTs 2.3.1), using brain-extracted versions of both T1w reference and the T1w template. The following templates were were selected for spatial normalization and accessed with *TemplateFlow* [23.0.0, @templateflow]: *FSL's MNI ICBM 152 non-linear 6th Generation Asymmetric Average Brain Stereotaxic Registration Model* [@mni152nlin6asym, RRID:SCR_002823; TemplateFlow ID: MNI152NLin6Asym], *ICBM 152 Nonlinear Asymmetrical template version 2009c* [@mni152nlin2009casym, RRID:SCR_008796; TemplateFlow ID: MNI152NLin2009cAsym].

#### Functional data preprocessing

For each of the 8 BOLD runs found per subject (across all tasks and sessions), the following preprocessing was performed. First, a reference volume and its skull-stripped version were generated by aligning and averaging 1 single-band references (SBRefs). Head-motion parameters with respect to the BOLD reference (transformation matrices, and six corresponding rotation and translation parameters) are estimated before any spatiotemporal filtering using `mcflirt` [FSL 6.0.3:b862cdd5, @mcflirt]. The BOLD time-series (including slice-timing correction when applied) were resampled onto their original, native space by applying the transforms to correct for head-motion. These resampled BOLD time-series will be referred to as *preprocessed BOLD in original space*, or just *preprocessed BOLD*. The

BOLD reference was then co-registered to the T1w reference using `mri_coreg` (FreeSurfer) followed by `flirt` [FSL 6.0.3:b862cdd5, @flirt] with the boundary-based registration [@bbr] cost-function.

Co-registration was configured with six degrees of freedom. First, a reference volume and its skull-stripped version were generated using a custom methodology of *fMRIPrep*. Several confounding time-series were calculated based on the *preprocessed BOLD*: framewise displacement (FD), DVARS and three region-wise global signals. FD was computed using two formulations following Power (absolute sum of relative motions, @power_fd_dvars) and Jenkinson (relative root mean square displacement between affines, @mcflirt). FD and DVARS are calculated for each functional run, both using their implementations in *Nipype* [following the definitions by @power_fd_dvars]. The three global signals are extracted within the CSF, the WM, and the whole-brain masks. Additionally, a set of physiological regressors were extracted to allow for component-based noise correction [*CompCor*, @compcor]. Principal components are estimated after high-pass filtering the *preprocessed BOLD* time-series (using a discrete cosine filter with 128s cut-off) for the two *CompCor* variants: temporal (tCompCor) and anatomical (aCompCor). tCompCor components are then calculated from the top 2% variable voxels within the brain mask. For aCompCor, three probabilistic masks (CSF, WM and combined CSF+WM) are generated in anatomical space. The implementation differs from that of Behzadi et al. in that instead of eroding the masks by 2 pixels on BOLD space, a mask of pixels that likely contain a volume fraction of GM is subtracted from the aCompCor masks. This mask is obtained by thresholding the corresponding partial volume map at 0.05, and it ensures components are not extracted from voxels containing a minimal fraction of GM.

Finally, these masks are resampled into BOLD space and binarized by thresholding at 0.99 (as in the original implementation). Components are also calculated separately within the WM and CSF masks. For each CompCor decomposition, the *k* components with the largest singular values are retained, such that the retained components' time series are sufficient to explain 50 percent of variance across the nuisance mask (CSF, WM, combined, or temporal). The remaining components are dropped from consideration. The head-motion estimates calculated in the correction step were also placed within the corresponding confounds file. The confound time series derived from head motion estimates and global signals were expanded with the inclusion of temporal derivatives and quadratic terms for each [@confounds_satterthwaite_2013]. Frames that exceeded a threshold of 0.5 mm FD or 1.5

standardized DVARS were annotated as motion outliers. Additional nuisance timeseries are calculated by means of principal components analysis of the signal found within a thin band (*crown*) of voxels around the edge of the brain, as proposed by [@patriat_improved_2017].

The BOLD time-series were resampled into several standard spaces, correspondingly generating the following *spatially-normalized, preprocessed BOLD runs*: MNI152NLin6Asym, MNI152NLin2009cAsym. First, a reference volume and its skull-stripped version were generated using a custom methodology of *fMRIPrep*. All resamplings can be performed with *a single interpolation step* by composing all the pertinent transformations (i.e. head-motion transform matrices, susceptibility distortion correction when available, and co-registrations to anatomical and output spaces). Gridded (volumetric) resamplings were performed using `antsApplyTransforms` (ANTs), configured with Lanczos interpolation to minimize the smoothing effects of other kernels [@lanczos]. Non-gridded (surface) resamplings were performed using `mri_vol2surf` (FreeSurfer).

Many internal operations of *fMRIPrep* use *Nilearn* 0.10.0 [@nilearn, RRID:SCR_001362], mostly within the functional processing workflow. For more details of the pipeline, see [the section corresponding to workflows in *fMRIPrep*'s documentation](https://fmriprep.readthedocs.io/en/latest/workflows.html "FMRIPrep's documentation").

### 3. Partial Least Squares analyses on Instruction period time series

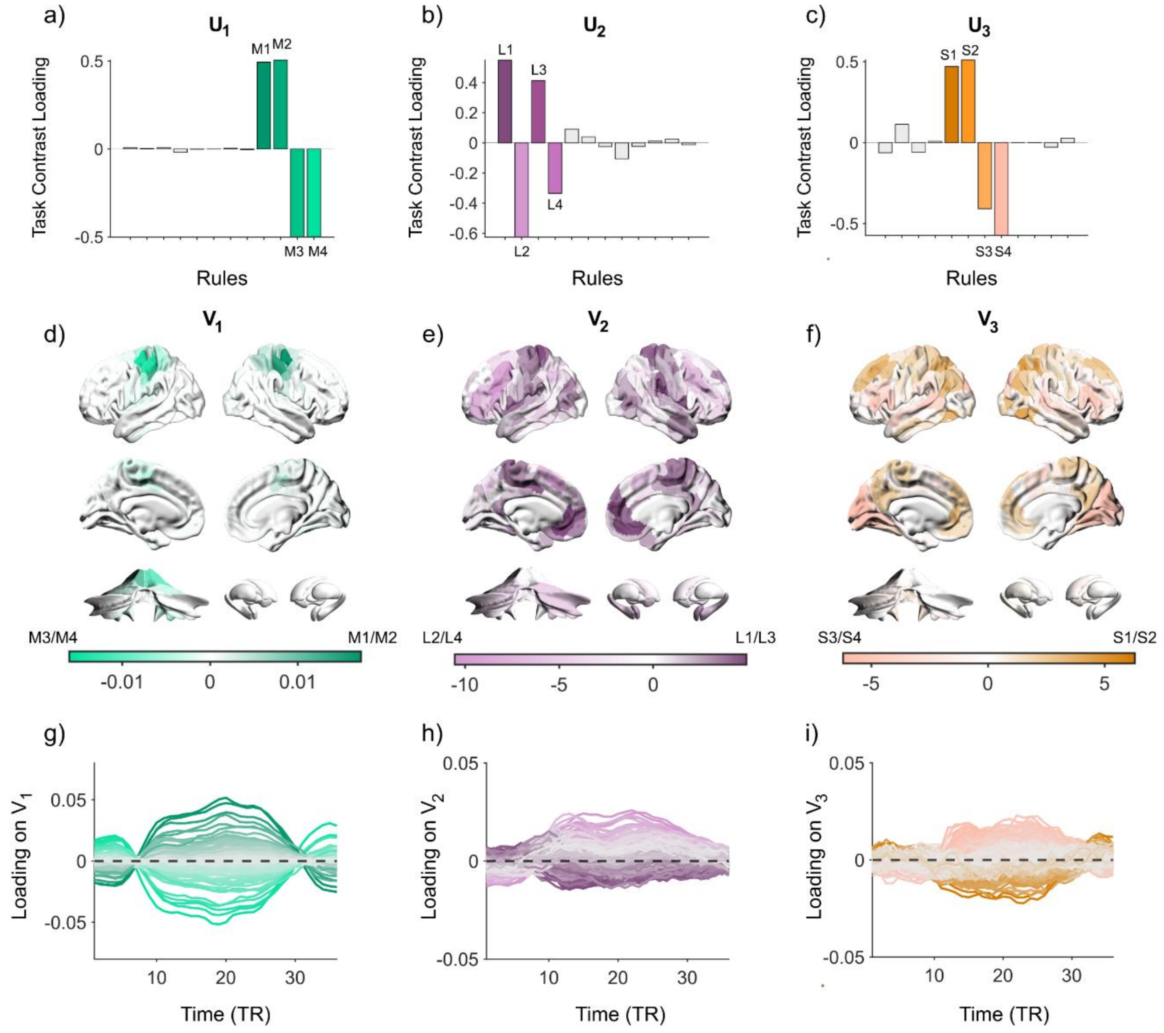


**Figure S1. Partial Least Squares Analysis on Instruction period time series.** a-c) Bar plot of the first 3 LVs of Task Contrast scores in PLS space. d-f) Brain visualisation of first 3 LVs of average brain activity in PLS space. g-i) Loadings of brain activity across time in PLS space. Each line denotes an individual region's dynamics across time.

### 4. Comparing Task Variance vs. Rule Bias in fMRI

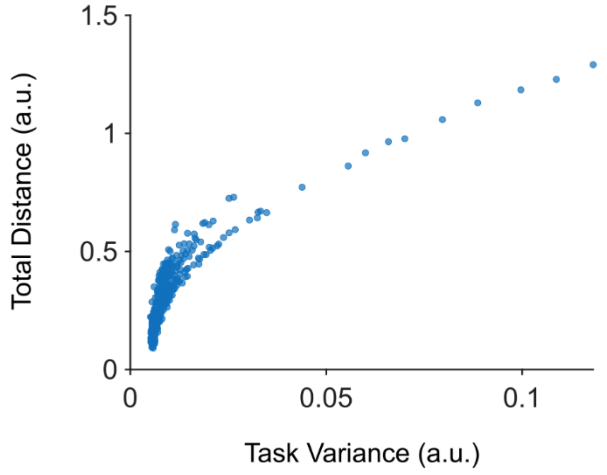


**Figure S2.** Scatter plot comparing task variance (measure from Yang et al., 2019) against rule bias (Euclidean distance in PLS space). Each dot is a brain region.

### 5. Table 1. Labels for Component brain regions

| ROI Name | X | Y | Z |
|---|---|---|---|
| 17Networks_LH_SomMotA_1 | -8 | -15 | 47 |
| 17Networks_LH_SomMotA_3 | -49 | -17 | 54 |
| 17Networks_LH_SomMotA_4 | -48 | -29 | 58 |
| 17Networks_LH_SomMotA_5 | -39 | -25 | 53 |
| 17Networks_LH_SomMotA_9 | -36 | -19 | 65 |
| 17Networks_LH_SomMotA_10 | -32 | -29 | 63 |
| 17Networks_LH_SomMotA_11 | -30 | -38 | 65 |
| 17Networks_LH_SomMotA_12 | -23 | -11 | 65 |
| 17Networks_LH_DorsAttnA_SPL_1 | -26 | -70 | 31 |
| 17Networks_LH_DorsAttnB_PostC_2 | -55 | -20 | 41 |
| 17Networks_LH_DorsAttnB_PostC_4 | -46 | -29 | 44 |
| 17Networks_LH_DorsAttnB_PostC_5 | -39 | -37 | 49 |
| 17Networks_LH_DorsAttnB_PostC_6 | -30 | -46 | 63 |
| 17Networks_LH_SalVentAttnA_FrOper_2 | -52 | 9 | 13 |
| 17Networks_LH_SalVentAttnB_PFCl_1 | -38 | 49 | 11 |
| 17Networks_LH_SalVentAttnB_Ins_2 | -33 | 25 | -1 |
| 17Networks_LH_ContA_IPS_1 | -29 | -74 | 42 |
| 17Networks_LH_ContA_IPS_3 | -35 | -62 | 48 |
| 17Networks_LH_ContA_PFClv_1 | -48 | 35 | 10 |
| 17Networks_LH_ContA_PFClv_2 | -42 | 38 | 22 |
| 17Networks_LH_ContA_PFCl_2 | -45 | 20 | 27 |
| 17Networks_LH_ContA_PFCl_3 | -39 | 7 | 34 |
| 17Networks_LH_ContB_IPL_3 | -42 | -52 | 49 |
| 17Networks_LH_ContB_PFCd_1 | -30 | 14 | 57 |
| 17Networks_LH_ContB_PFClv_1 | -42 | 49 | -6 |
| 17Networks_LH_ContB_PFCmp_1 | -4 | 28 | 47 |
| 17Networks_LH_DefaultB_PFCd_6 | -6 | 10 | 65 |
| 17Networks_LH_DefaultB_PFCl_1 | -41 | 19 | 48 |
| 17Networks_LH_DefaultB_PFCl_2 | -42 | 7 | 48 |
| 17Networks_LH_DefaultB_PFCv_4 | -48 | 28 | 0 |
| 17Networks_LH_DefaultB_PFCv_5 | -53 | 19 | 11 |
| 17Networks_RH_SomMotA_1 | 54 | -17 | 40 |
| 17Networks_RH_SomMotA_2 | 52 | -13 | 49 |
| 17Networks_RH_SomMotA_4 | 49 | -26 | 56 |
| 17Networks_RH_SomMotA_5 | 7 | -10 | 51 |

| | | | |
|---|---|---|---|
| 17Networks_RH_SomMotA_6 | 43 | -21 | 54 |
| 17Networks_RH_SomMotA_7 | 37 | -20 | 64 |
| 17Networks_RH_SomMotA_9 | 31 | -41 | 64 |
| 17Networks_RH_SomMotA_10 | 34 | -27 | 61 |
| 17Networks_RH_SomMotA_12 | 29 | -11 | 65 |
| 17Networks_RH_DorsAttnA_TempOcc_3 | 50 | -64 | -9 |
| 17Networks_RH_DorsAttnA_SPL_2 | 32 | -66 | 35 |
| 17Networks_RH_DorsAttnA_SPL_6 | 34 | -50 | 54 |
| 17Networks_RH_DorsAttnB_PostC_3 | 44 | -37 | 50 |
| 17Networks_RH_DorsAttnB_PostC_4 | 45 | -28 | 42 |
| 17Networks_RH_DorsAttnB_PostC_5 | 35 | -36 | 51 |
| 17Networks_RH_DorsAttnB_PostC_7 | 24 | -50 | 68 |
| 17Networks_RH_SalVentAttnB_Ins_2 | 37 | 23 | 5 |
| 17Networks_RH_ContA_PFCl_2 | 48 | 18 | 23 |
| 17Networks_RH_ContA_PFCl_3 | 47 | 29 | 28 |
| 17Networks_RH_ContB_PFCmp_1 | 5 | 28 | 48 |
| 17Networks_RH_DefaultB_PFCv_3 | 54 | 24 | 6 |
| Left_V | -18 | -50 | -19 |
| Right_V | 18 | -50 | -19 |

#### 6. Table 2. Labels for Recombination brain regions

| ROI Name | X | Y | Z |
|---|---|---|---|
| 17Networks_LH_SomMotB_Aud_1 | -50 | -9 | 0 |
| 17Networks_LH_SomMotB_Aud_2 | -56 | -22 | 8 |
| 17Networks_LH_SomMotB_Ins_1 | -36 | -24 | 10 |
| 17Networks_LH_SomMotB_Aud_3 | -59 | -37 | 16 |
| 17Networks_LH_SomMotB_Aud_4 | -41 | -35 | 14 |
| 17Networks_LH_DorsAttnA_SPL_4 | -29 | -58 | 50 |
| 17Networks_LH_DorsAttnB_PostC_3 | -55 | -32 | 45 |
| 17Networks_LH_DorsAttnB_FEF_1 | -40 | -3 | 51 |
| 17Networks_LH_DorsAttnB_FEF_2 | -25 | -1 | 55 |
| 17Networks_LH_DorsAttnB_FEF_3 | -30 | -8 | 52 |
| 17Networks_LH_DorsAttnB_PrCv_1 | -50 | 3 | 38 |
| 17Networks_LH_SalVentAttnA_ParOper_1 | -55 | -32 | 22 |
| 17Networks_LH_SalVentAttnA_ParOper_2 | -58 | -44 | 27 |
| 17Networks_LH_SalVentAttnA_Ins_3 | -33 | 19 | 8 |
| 17Networks_LH_SalVentAttnA_FrMed_2 | -5 | 9 | 48 |
| 17Networks_LH_ContA_IPS_2 | -58 | -42 | 45 |
| 17Networks_LH_ContA_IPS_4 | -45 | -41 | 47 |
| 17Networks_LH_ContA_IPS_5 | -33 | -46 | 41 |
| 17Networks_LH_ContA_PFCl_1 | -49 | 6 | 26 |
| 17Networks_LH_TempPar_3 | -62 | -32 | 5 |
| 17Networks_LH_TempPar_4 | -52 | -43 | 5 |
| 17Networks_LH_TempPar_6 | -59 | -49 | 16 |
| 17Networks_RH_SomMotB_Aud_1 | 53 | 3 | -6 |
| 17Networks_RH_SomMotB_Aud_2 | 53 | -14 | 6 |
| 17Networks_RH_SomMotB_Ins_1 | 39 | -19 | 5 |
| 17Networks_RH_SomMotB_Aud_3 | 60 | -24 | 11 |
| 17Networks_RH_SomMotB_S2_4 | 41 | -29 | 18 |
| 17Networks_RH_DorsAttnB_FEF_1 | 39 | -3 | 53 |
| 17Networks_RH_SalVentAttnA_PrC_1 | 51 | 3 | 41 |
| 17Networks_RH_SalVentAttnA_FrOper_3 | 54 | 12 | 12 |
| 17Networks_RH_SalVentAttnA_FrMed_2 | 6 | 11 | 58 |
| 17Networks_RH_SalVentAttnA_FrMed_3 | 7 | -2 | 67 |
| 17Networks_RH_SalVentAttnA_FrMed_4 | 16 | 7 | 69 |
| 17Networks_RH_SalVentAttnB_PFCl_1 | 42 | 46 | 14 |
| 17Networks_RH_ContA_IPS_2 | 54 | -33 | 51 |
| 17Networks_RH_ContA_IPS_3 | 47 | -44 | 46 |
| 17Networks_RH_ContA_IPS_4 | 36 | -44 | 45 |
| 17Networks_RH_ContA_PFCl_4 | 49 | 8 | 25 |
| 17Networks_RH_ContA_PFCl_5 | 39 | 11 | 34 |

| | | | |
|---|---|---|---|
| 17Networks_RH_ContB_IPL_3 | 56 | -41 | 48 |
| 17Networks_RH_ContB_IPL_4 | 41 | -55 | 48 |
| 17Networks_RH_ContB_PFCld_1 | 39 | 33 | 38 |
| 17Networks_RH_ContB_PFCld_2 | 45 | 19 | 44 |
| 17Networks_RH_ContB_PFCld_3 | 43 | 7 | 51 |
| 17Networks_RH_TempPar_6 | 59 | -46 | 7 |
| 17Networks_RH_TempPar_8 | 65 | -34 | 11 |
| Left_VI | -30 | -50 | -27 |
| Right_VI | 30 | -50 | -27 |
| Left_VIIb | -33 | -65 | -53 |
| Right_VIIb | 33 | -65 | -53 |
| Right_VIIIa | 20 | -65 | -53 |

### 7. Delta Functional Connectivity Matrices

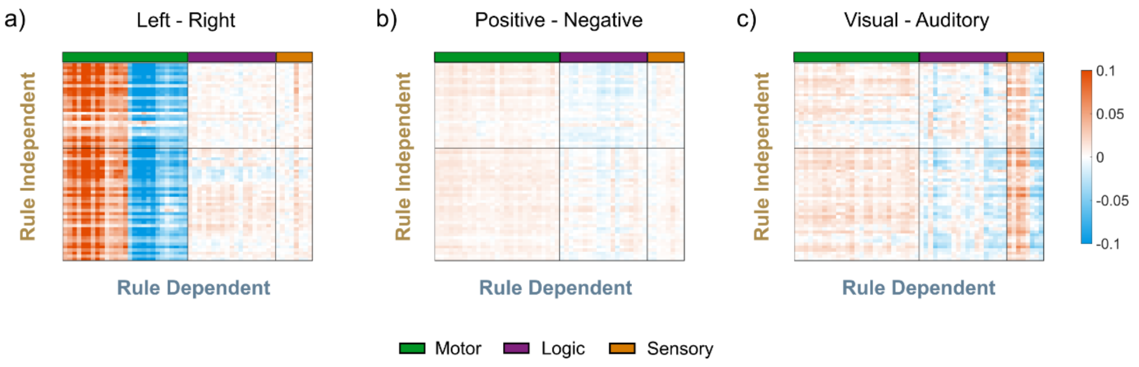


**Figure S3. Average difference in functional connectivity between rule dependent and rule independent regions for each domain.** a) Motor domain: left hand – right hand. b) Logic domain: positive – negative. c) Sensory domain: visual – auditory. Rule dependent regions (columns) were assigned to specific domains (green = motor, purple = logic, orange = sensory) based off their strongest loading on the PLS axes.

### 8. Recombination and Component characteristics with strict Bootstrap threshold

To assess whether characteristics of Recombination regions were driven by regions stable only on one latent variable, we applied a stricter criteria, including only regions that were stable on all three latent variables (|BSR| > 2 on three latent variables). Overall, 46/51 Recombination and 51/54 Component regions passed this threshold (Figure S4a/b).

Dimensionality between Recombination and Component regions were compared using a paired t-test. Component regions had significantly higher dimensionality compared to Recombination regions ($t_{86}$ = 10.69, p < 1e-16, 95% CI [1.16 1.69]; Figure S4c). Participation coefficient between the two groups were compared using paired t-test. Recombination regions were found to be significantly more integrated with the rest of the brain compared to Component regions ($t_{95}$ = 2.52, p = 0.01, 95% CI [0.002 0.015]; Figure S4d). Recombination regions were more correlated than Component regions across mini-blocks ($t_{8,437}$ = 42.99, p < 1e-16, 95% CI [0.12 0.13]; Figure S4e). Therefore, Recombination regions were low-dimensional, integrated, and generalised across task contexts, replicating our main findings.

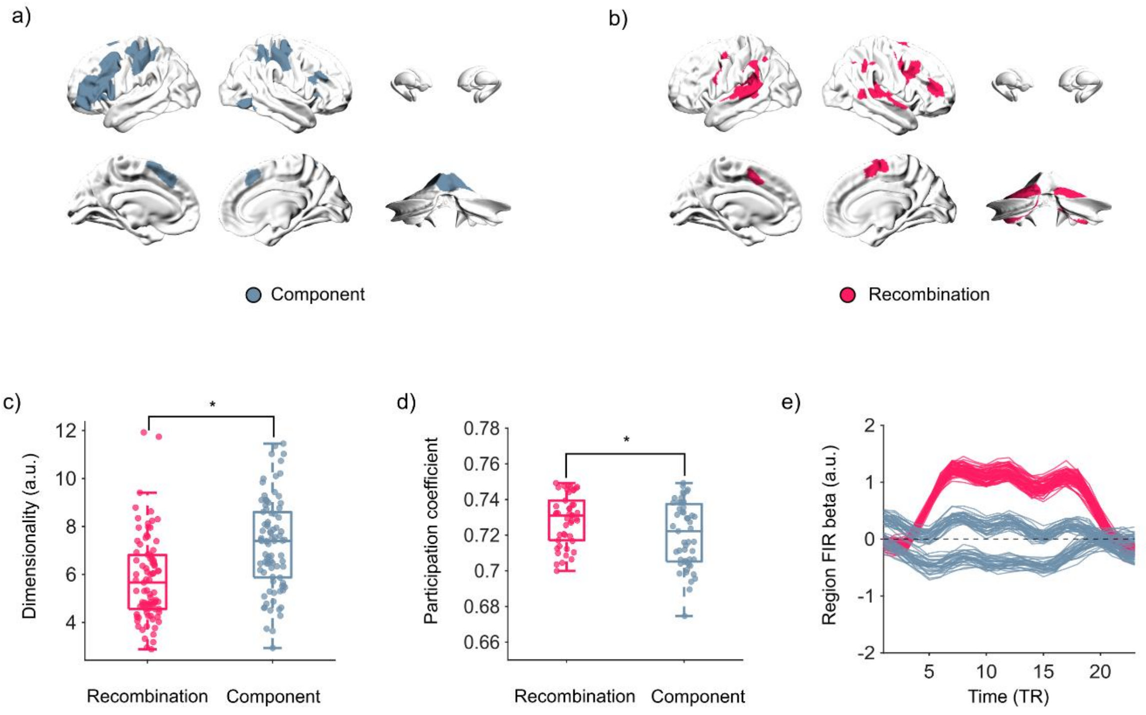


**Figure S4. Characteristics of Recombination regions using strict Bootstrap threshold (|BSR| > 2 on all three latent variables)**. a) Brain visualisation of Component regions. b) Brain visualisation of Recombination regions. c) Boxplot of Recombination (pink) and Component (silver) regions dimensionality measured by the participation ratio. d) Boxplot of participation coefficient scores for Recombination (pink) and Component (silver) regions. e) Group average beta estimates across time for an example region from Recombination (pink)

and Component (silver) regions. Each line is the FIR estimate for a different mini-block. For boxplots in a) and b), each dot is a brain region. Centre line, median, box limits, upper and lower quartiles. Whiskers, 1.5x interquartile range. * denotes significant difference $p < 0.05$.
